# msallsearchpy: Universal Search Engine with Enhanced Performance for High-Flow Quantitative LC-MS Proteomics

**DOI:** 10.64898/2026.10.16.759407

**Authors:** Mark V. Ivanov, Leyla A. Garibova, Daria D. Emekeeva, Valeriy I. Postoenko, Elizaveta M. Kazakova, Andrey A. Shelepchikov, Alexey V. Tretyakov, Irina A. Tarasova, Mikhail V. Gorshkov

## Abstract

Recent studies have demonstrated that the flow rates of a few µl/min can be a more efficient alternative to nanoLC-based proteome analysis. We present a novel search engine, msallsearchpy, applicable to data acquired using all existing proteome-wide acquisition methods including DIA, DDA and MS1-only, with enhanced efficiency for higher flow rate analysis. For the data acquired using Orbitrap FTMS at the flow rates of up to 400 µl/min, msallsearchpy demonstrates significantly improved identification and quantification performance compared with the state-of-the-art DDA and DIA alternatives. As one of the striking examples, msallsearchpy provides 2000 protein identifications for the DDA data acquired for 20 ng proteomic samples and flow rate of 50 µL/min, compared with the none by popular search engines. The results provide further support to the currently floating ideas in quantitative proteomics regarding the use of high-flow rates as a more efficient alternative to nanoLC.

## INTRODUCTION

The idea to create a universal search pipeline for all kinds of proteomics data is not new. A recent example is CHIMERYS, a spectrum-centric search engine that unifies data-dependent (DDA), data-independent (DIA) and parallel reaction monitoring (PRM) data analyses^1^. Another example is the FragPipe computational platform which provides workflows for both DDA and DIA methods^2^. Previously, we showed that despite the lower number of total identifications, MS1-only proteome-wide analysis is capable of providing quantification efficiency comparable with the standard DDA/DIA methods^3,4^. This comparison is even more pronounced under short HPLC gradient conditions^5,6^. In the core of MS1-only method called DirectMS1 is a search engine ms1searchpy^7^ and peptide feature detection utility Biosaur^8^, which allow extraction of peptide isotopic ion envelopes directly from MS1 spectra using all available complementary information about peptides for the subsequent protein identifications.

In this work we modified the ms1searchpy search engine to make it capable for utilizing MS/MS information as well when the proteome analysis was performed using either DDA, or DIA approaches, thus transforming it to a universal search engine for all types of large scale proteome characterization methods. Although the versatility of the search engine itself in terms of data collection methods is not a mandatory feature, the biggest novel and unique advantage of the presented development is its efficiency for high-flow rate applications. Indeed, the idea to move away from nanoflow proteomics to higher flow rates has been actively discussed recently^9–13^. However, the price for further flow rate increase is the large sample consumption at the levels prohibited for many applications. The overcoming of this limitation was successfully demonstrated in a number of works showing identification of thousands of proteins in short LC gradient analyses using analytical flow rate chromatography and low sample loading amounts. For example, the Scanning SWATH method has reported 2000 proteins identified for 250 ng of tryptic digest analysed at 800 ul/min flow rate with 5 min LC gradient^14^. Comparable results were obtained later using ZenoTOF and timsTOF Pro instruments and micro- and/or analytical-flow rate chromatography^15,16^. Notably, all these previous demonstrations were performed using time-of-flight (TOF) mass analyzers. Attempts to perform proteome wide analyses using higher flow rates and low sample amounts on the high-resolution Orbitrap instruments and employing existing state-of-the art MS/MS-based search engines have been less impressive demonstrating meaningful efficiency at flow rates of a few µl per minutes at best^10,11,13^. Our premise and the motivation for this work are that the reason for this discrepancy in results lies in the fundamental approach to identification used in existing search algorithms, which rely exclusively on fragmentation spectra. The search engine developed in this work overcomes this limitation demonstrating its superior capabilities in proteome analyses of sample in minute quantities performed using high-resolution Orbitrap FTMS and high-flow rate separations.

## RESULTS

### MS1-based search algorithm with MS/MS spectra processing extension provides a universal solution for all types of data acquisition methods

The developed search algorithm, msallsearchpy, performs identification of proteins from peptide isotopic envelopes (peptide features) in MS1 spectra, similar to the one of DirectMS1 method, with additional assistance from MS/MS spectra. With the latter assistance, some of the peptide-feature matches (PFM) found in MS1 spectra are supplemented with available fragmentation spectra. The workflow implemented in the search algorithm and shown in Figure 1 is described in more detail in the Methods section.

**Figure 1.**
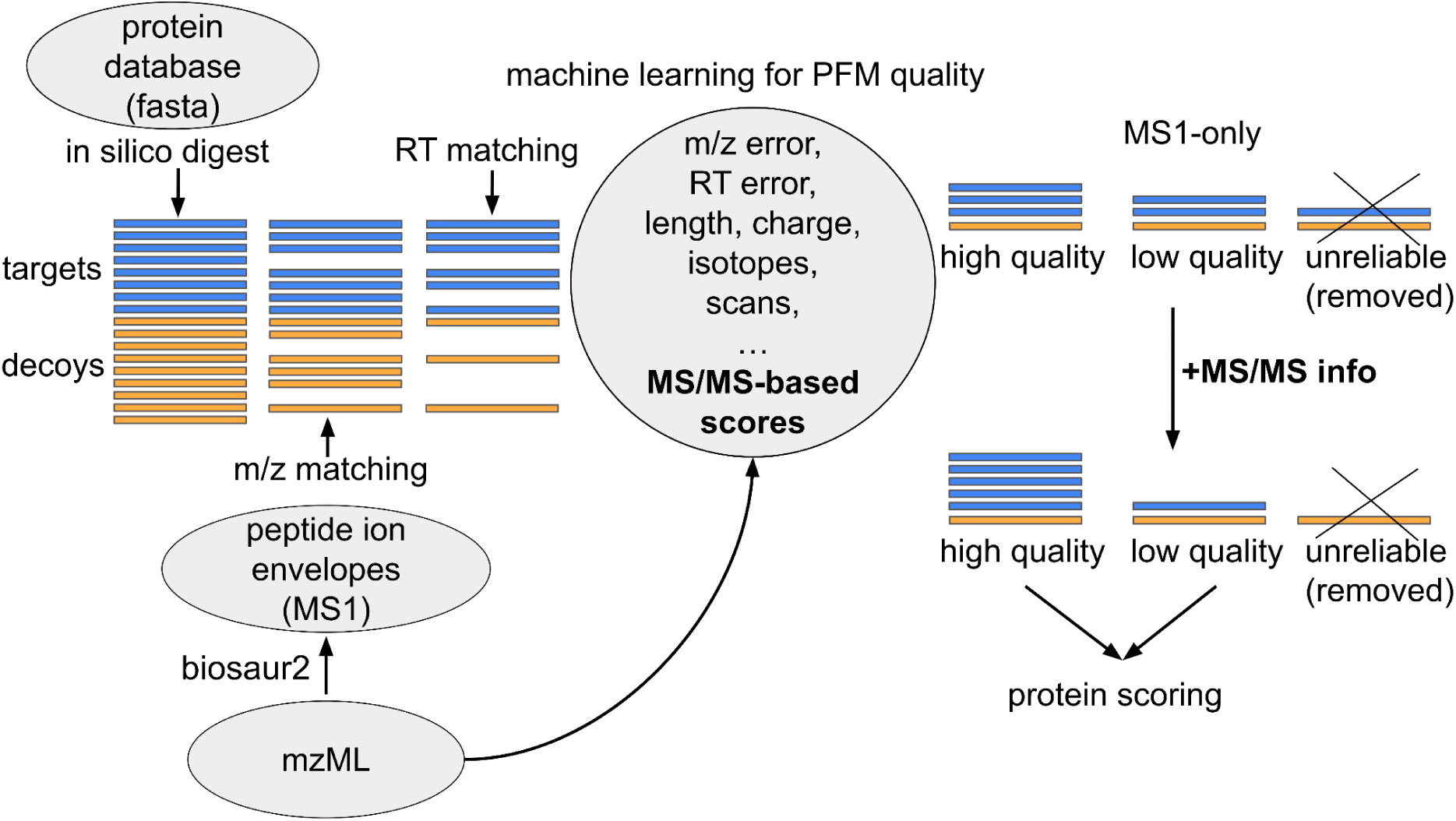
msallsearchpy workflow design. It starts with untargeted detection of peptide isotopic envelopes in experimental MS1 spectra using biosaur2 algorithm. Next, the msallsearchpy digests proteins *in silico* from the search database to obtain the list of theoretical peptides, followed by matching the peptide features with theoretical peptides using theoretical masses and predicted retention times (RT). At the machine learning-based classification stage, the PFM descriptors in ms1searchpy algorithm are supplemented with scores calculated from the MS/MS spectral matches of fragmented PFMs. The final step of the workflow is the same protein scoring implemented in the DirectMS1 method.

Supplementary Figure S1 illustrates how the two key steps in the msallsearchpy, RT prediction and MS/MS spectra analysis, affect the number of identified protein groups at 1% FDR. These results were obtained using DIA data acquired on an Orbitrap QE HF mass spectrometer with the MS1 resolution set to 60k. Both steps were evaluated for three possible implementations. They were “no RT prediction”, additive model based on retention coefficients, and machine-learning-based DeepLC^17^ for RT prediction, and “no fragmentation”, simple calculation of fragment matches, and machine-leaning-based fragmentation pattern prediction using MS2PIP^18^. Expectedly, when MS/MS spectra are not utilized for the search, retention time prediction becomes crucially important as the only source of peptide’s sequence-specific information. DeepLC is the obvious winner in this race because the additive RT prediction model is not sequence-specific by its nature^19^. MS/MS spectra add sequence-specificity that results in significant improvement in the search efficiency. Surprisingly, the particular implementation of theoretical MS/MS spectra construction is not important, and a straightforward fragment m/z matching produces results similar to the ones with fragmentation intensities prediction. RT prediction is still important, yet the effect is less profound. Supplementary Figure S2 illustrates further the distribution of target and decoy PFMs by their respective quality metrics in the MS1-only and MS1+MS/MS assisted analyses, as well as by the numbers of b and y ions found. Based on the above results, the rest of the study was performed using DeepLC for RT prediction and simple matched fragment calculations for MS/MS spectra analysis.

### High-flow DDA public datasets

The recent work by Tsiklauri et al. highlights the growing shift from nanoflow to microflow separation in modern proteomic workflows^10^. Notably, the authors demonstrated that capillary LC at a flow rate of 1.5 μL/min achieves protein identification efficiency comparable to nanoflow LC across a wide range of sample loads of 5 to 10,000 ng, while significantly improving quantitative reproducibility. Furthermore, the study showed that increasing the flow rate to 50 μL/min provided the highest quantitative reproducibility.

However, this benefit was limited at low sample loads, and the standard search engines failed to yield a meaningful number of protein identifications for the sample amounts below 500 ng. To compare our algorithm’s performance with standard search engines, we re-analysed this data using msallsearchpy, with the results presented in Figure 2a. In particular, the results obtained using IdentiPy search engine^20^, developed and optimized for the analysis of DDA data, were expected: at high-flow rates the number of identifications for low sample amounts decreased. For the large amounts of sample loads (2000 ng), there were no differences in the search efficiencies observed in a wide range of flow rates. In contrast, the MS1-based search yielded the opposite results demonstrating that the 50 μL/min flow rate remained consistently robust across all sample amounts. Supplementary Figure S3 provides a validation of these findings, showing an extremely high degree of overlap (ca.98%) between the proteins reported by msallsearchpy at a 20 ng sample load and those identified by IdentiPy for 2000 ng load data. Furthermore, we performed multi-species searches using a combined human, *E. coli*, and yeast protein database (Supplementary Figure S4). Based on the relative database size, the expected proportion of false entrapment identifications (*E. coli* and yeast) was 0.35% of total IDs at 1% FDR. The empirical false discovery proportion (FDP) estimated using the “combined” method^21^ ranged from 0.5% to 2%, with an average of 1.25%. The “combined” method provides an estimated upper bound of FDP and on average it overestimates the true FDR, thus, explaining the deviation of the average from 1%. Notably, the choice of standard DDA search engine did not affect the results, as demonstrated by direct comparison between IdentiPy and the FragPipe (Supplementary Figure S5).

**Figure 2.**
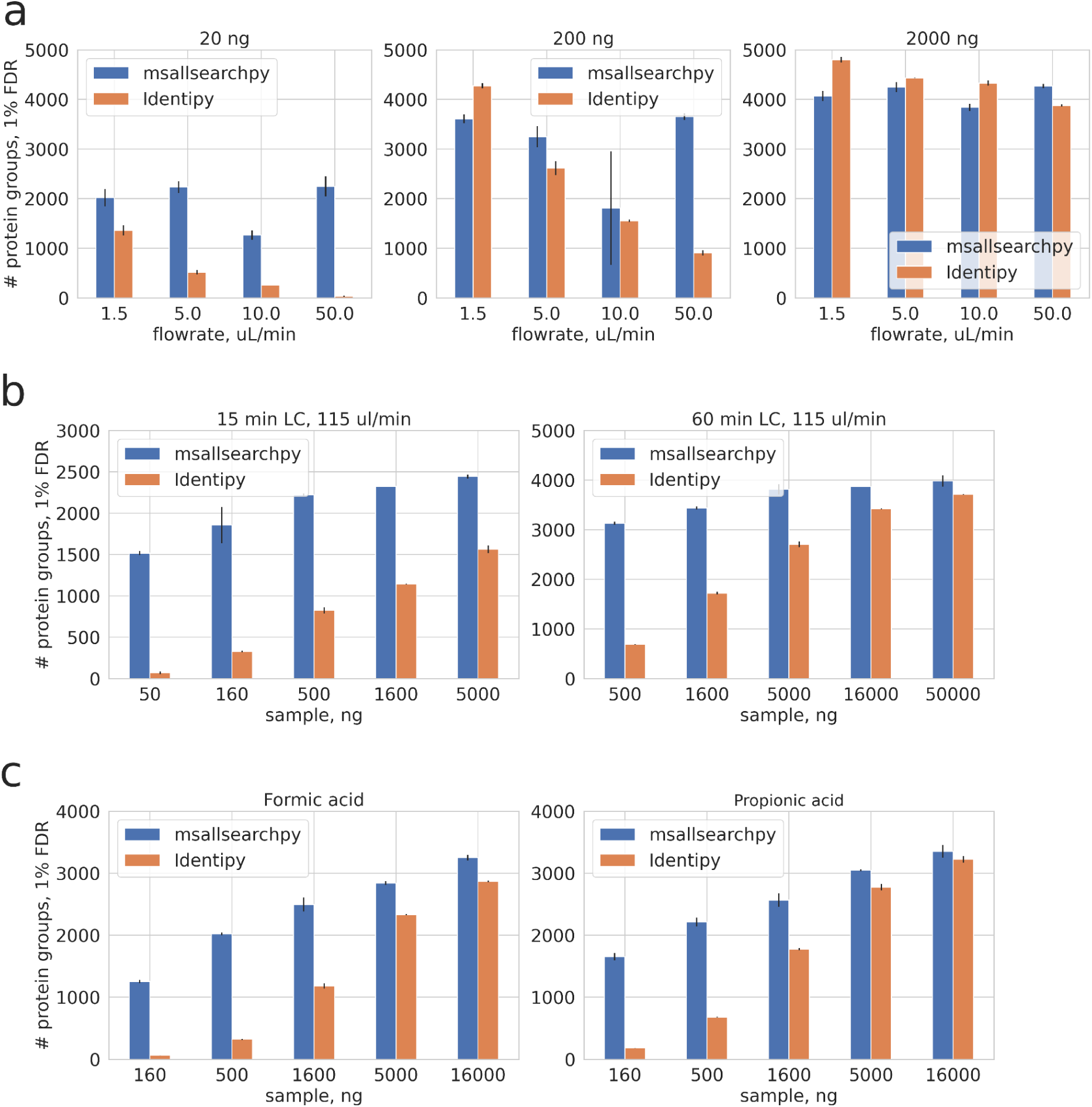
The number of identified protein groups at 1% FDR for different high-flow DDA datasets: (a) Tsiklauri et al. (b) Jadeja et al. (c) Starovoit et al. Results for msallsearchpy are indicated in blue, while results for the traditional DDA search engine, Idenipy, are shown in orange.

To further explore the performance of msallsearchpy in high-flow rate proteomics, we re-analysed the data from Jadeja et al.^13^, which focused on comparing microbore columns with 1.5 and 1.0 mm internal diameters for microflow applications. Our reanalysis, presented in Figure 2b demonstrates that the developed search engine consistently outperforms alternative algorithms at high-flow rates, regardless of the gradient duration or sample load. Furthermore, the results obtained for the 1.0 mm i.d. column did not differ significantly those of the 1.5 mm i.d. column and are provided in Supplementary Figure S6.

Next, we re-analysed data from Starovoit et al.^9^ acquired under microflow conditions for different sample loads and two different mobile phase acidifiers, formic and propionic acids (Figure 2c). Importantly, the results not only confirm the unique performance of msallsearchpy but also support the conclusion from the referenced study regarding the potential of utilizing propionic acid in bottom-up high-flow rate proteomics.

In summary, our results demonstrate the superior performance of the MS1-centric search concept compared to traditional MS/MS-centric approaches in high-flow rate proteomic analyses. The developed msallsearchpy engine is particularly well-suited for high throughput proteomics involving minute sample amounts at elevated flow rates, with the latter providing enhanced reproducibility and quantification accuracy.

### Quantitative datasets

Our findings highlight the specific advantages of msallsearchpy for the proteomic analysis of low sample amounts under the following conditions: high resolution mass-spectrometry (e.g. Orbitrap FTMS) with a high MS1 spectra acquisition rate to ensure sufficient number of spectra, analytical or microflow rates, wide-bore LC columns, DDA method, and short LC gradients. To further demonstrate these advantages we performed a quantitative proteome analysis of varying sample loads for ovarian cancer cell line A549 treated with Bortezomib. This analysis utilized the DIA method, 400 ul/min flow rate, and 20 min LC gradient as detailed in the Methods section. The FragPipe-DIA was used as the standard DIA search workflow for comparison with msallsearchpy (Figure 3a). At a 5000 ng sample load, the FragPipe outperformed msallsearchpy in the number of reported differentially expressed proteins (DEPs). However, when the sample load was reduced to 500 ng, FragPipe-DIA exhibited a dramatic decrease to only 5 DEPs compared to 101 DEPs reported by msallsearchpy. Note that 77 of the latter DEPs were also presented in the FragPipe’s DEPs obtained at the 5000 ng load.

**Figure 3.**
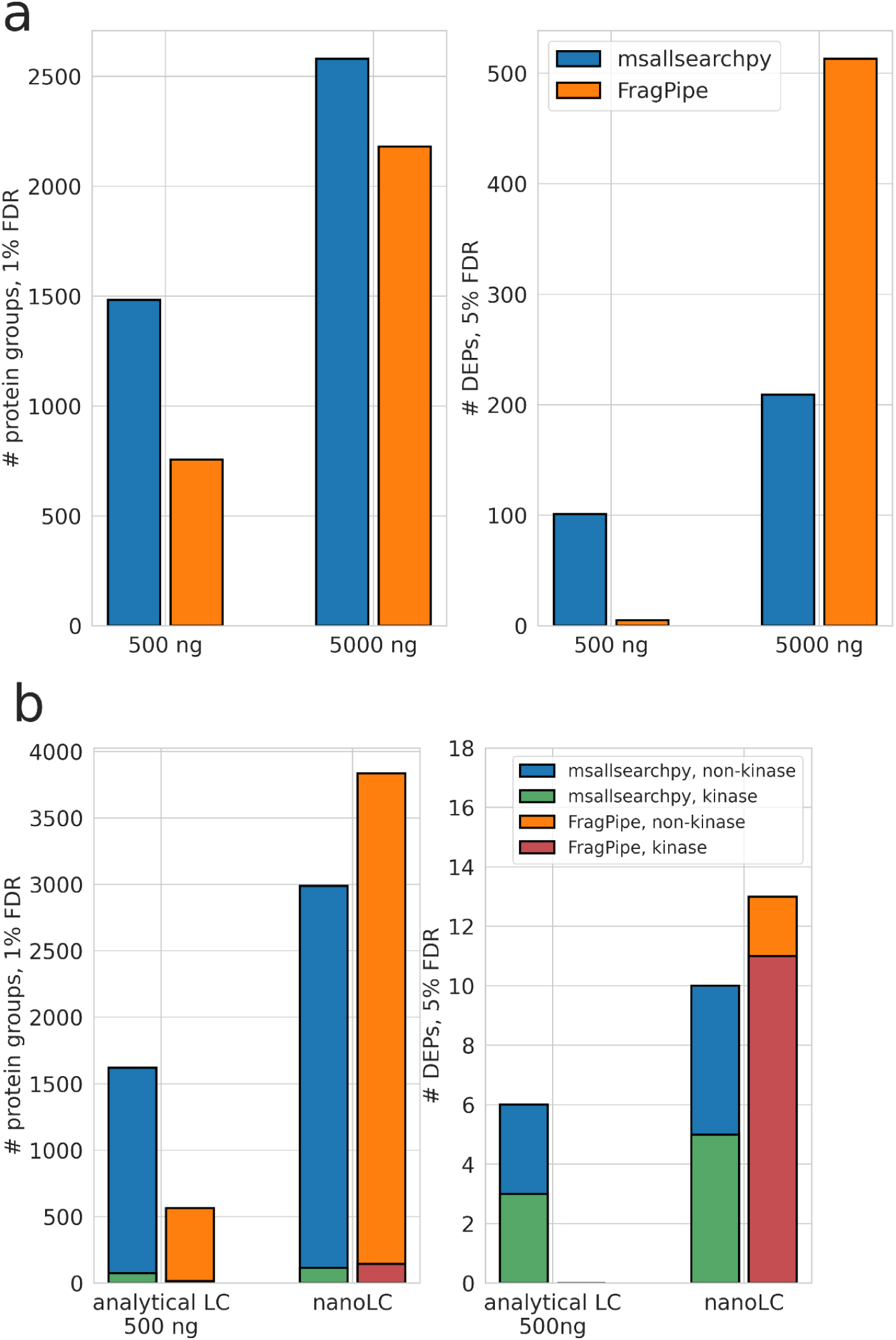
The number of identified protein groups and differentially expressed proteins reported by msallsearchpy+DirectMS1Quant and FragPipe+FragPipeAnalyst workflows. Results were filtered using 0.05 p-value threshold with Benjamini-Hochberg correction. Fold change thresholds were: 0.5 absolute log2 value for FragPipeAnalyst and automatic for DirectMS1Quant. (a) Results are shown for ovarian cancer cell line A549 treated with Bortezomib, DIA acquisition, 400 ul/min flow rate, 2.1 mm i.d. LC column and 20 min LC gradient. Results were filtered using 0.05 p-value threshold with Benjamini-Hochberg correction. Fold change thresholds were: 0.5 absolute log2 value for FragPipeAnalyst and automatic for DirectMS1Quant. (b) Analytical LC results are shown for cancer cell line HCT116 treated with Staurosporine, DIA acquisition, PISA (7 temperature points), 400 ul/min flow rate, 2.1 mm i.d. LC column and 20 min LC gradient. nanoLC results represent data from Batth et al.: DIA acquisition, PISA (3 temperature points), 250 nl/min flow rate and 50 min LC gradient.

We next performed a thermal proteome profiling (TPP) of HCT116 cell lysates treated with Staurosporine (and DMSO as a vehicle control) using protein solubility alteration assay (PISA) and the same analytical DIA setup (Figure 3b). FragPipe-DIA failed to report any kinase DEPs, which are the expected targets of Staurosporine, whereas msallsearchpy successfully reported three kinases. As a reference, we re-analysed data from Batth et al.^22^, where a similar sample preparation protocol based on centrifugation was utilized. The primary differences compared to our experiments include a 2.5-fold longer gradient, the use of nanoLC, and highly optimized temperature points for Staurosporine within the PISA assay (3 pooled points versus 7 points in our setup). For these nanoLC data, FragPipe outperformed msallsearchpy across all evaluated metrics.

### LFQ Bench Astral (NanoLC)

In the final round of evaluation, we processed DDA data from Puyvelde et al. obtained for a 250 pg mixture of *E. coli* and yeast digests at predefined relative concentrations. Data was acquired using Orbitrap Astral. msallsearchpy yielded a greater number of DEPs at 5% empirical quantitative FDR, a more accurate quantitative FDR estimation without prior knowledge of false DEPs, a higher partial Area Under the Curve (pAUC) up to 5% FDR, and an increased number of identified protein groups (Figure 4). Conversely, the FragPipe’s quantitation workflow exhibited superior accuracy and precision in fold change estimation. While the results for the DIA data from the same study are also provided in Supplementary Figure S7, their interpretation warrants caution due to technical considerations detailed in the Discussion section.

**Figure 4.**
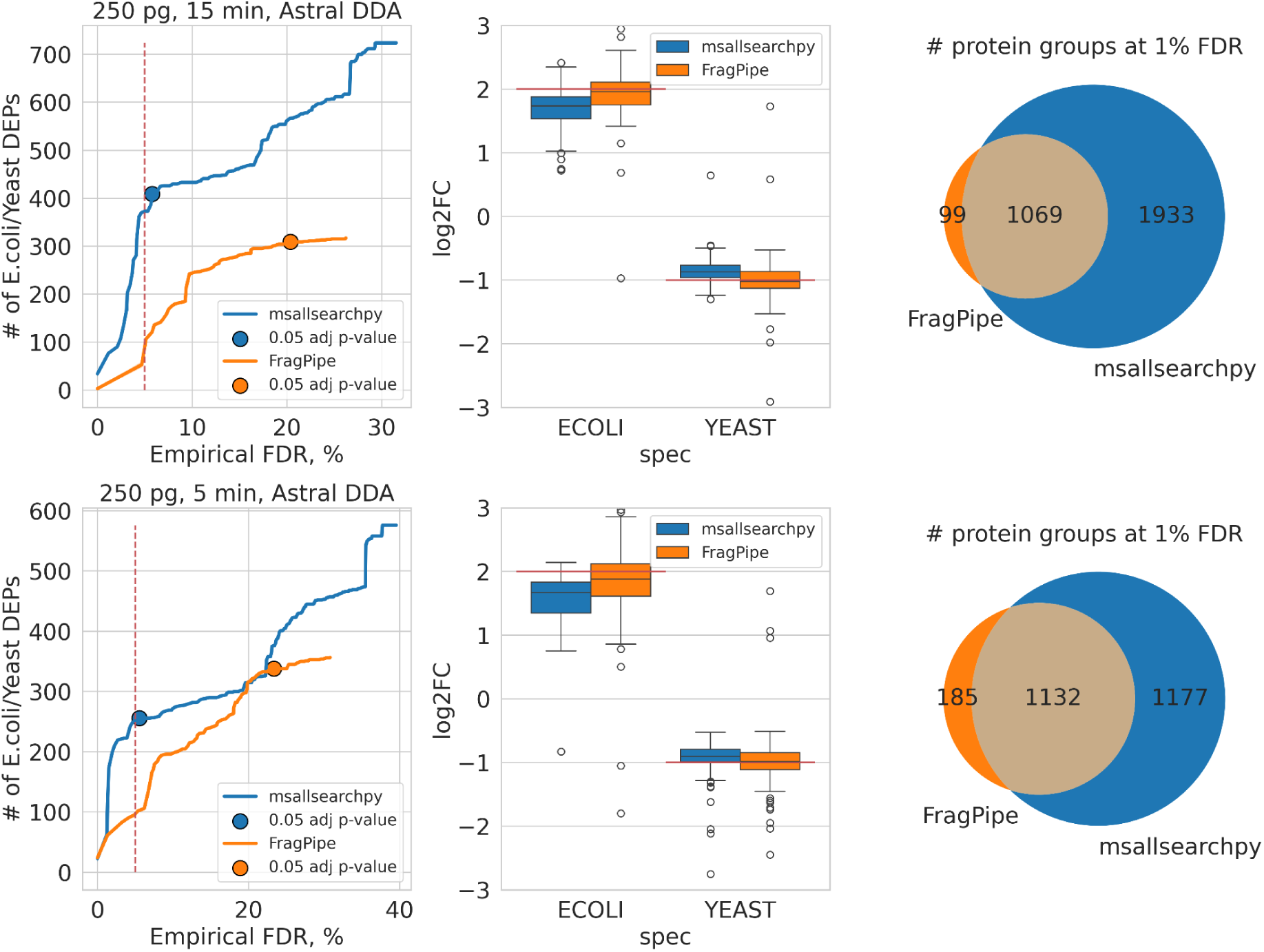
Results for 250 pg Orbitrap Astral DDA data from Puyvelde et al. analysed by msallsearchpy+DirectMS1Quant (blue) and FragPipe+FragPipeAnalyst (orange) workflows: (upper figures) 15 min LC and (bottom figures) 5 min LC data. (left column) The number of reported differentially expressed *E. coli* and yeast proteins. Empirical FDR was calculated using reported human DEPs. Only the proteins that passed fold change thresholds were used for the plots. Fold change thresholds were: 0.5 absolute log2 value for FragPipeAnalyst and automatic for DirectMS1Quant. Blue and orange dots represent the results which were obtained using 0.05 p-value threshold with Benjamini-Hochberg correction with no prior knowledge on the empirical FDR. *E. coli* and yeast proteins with opposite to the real changes sign of log2 fold changes were considered as false DEPs. (middle column) The log2 fold changes of DEPs. Red lines represent the experimental fold changes. (right column) The number of identified protein groups at 1% FDR.

## Discussion

The results in Figure 2 show that the msallsearchpy solves the problem of low identification efficiency of existing DDA search engines when it comes to the proteome analyses of low amounts of samples at high-flow rates. The latter is an especially attractive feature which has gained attention in proteomics recently due to its reproducibility, quantification accuracy and the ability to analyse thousands of samples in a row without disruptions due to the issues with chromatographic systems. The presented results reveal a greater potential for the recently proposed high-flow rate methods of proteomic analysis exceeding the expectations of their original developers. Our quantitative DIA data acquired at a 400 ul/min flow rate demonstrate that proteins reported by msallsearchpy are not merely identified but also robustly quantified with the workflow consistently outperforming standard search engines as sample loads decrease. We argue further that the number of differentially expressed proteins detected by a given workflow should serve as the primary benchmark for evaluating its quantification performance. Indeed, this metric directly reflects the method’s capability to resolve biological questions contrary to the widely employed coefficient of variation (CV) and the likes which are fundamentally limited when assessing the true quantitative power of a search pipeline^23^.

To demonstrate that msallsearchpy is applicable beyond high-flow rate analysis, we evaluated its performance using standard nanoLC experimental datasets. Our results, obtained for 250 pg sample loads using a state-of-art Orbitrap Astral mass-spectrometer in DDA mode, indicate that the search engine can effectively replace existing solutions for short gradients and low sample amounts. Several factors explain why the performance was markedly reduced under DIA acquisition. First, the DIA datasets contained a lower number of acquired MS1 spectra compared to DDA data (500 and 1600 spectra for 5 and 15 min DIA, vs. 2100 and 3700 spectra for 5 and 15min DDA, respectively). Second, the DIA acquisitions utilized FAIMS separation with a single compensation voltage (CV). While multiple FAIMS CV values are beneficial for MS1-centric acquisition [10.1021/acs.jproteome.0c00863], employing a single CV value limits the peptides’ detectability in MS1 spectra, consequently reducing protein sequence coverages and their subsequent identification scores. Finally, the m/z range of MS1 spectra acquisitions in the DIA was nearly half as wide as the m/z range utilized for DDA acquisition.

In recent years, numerous DIA method optimizations, including those provided by MS instrument vendors, have focused on maximizing the number of identified proteins when using popular DIA search engines such as DIA-NN or Spectronaut. However, these search engines do not rely on MS1-level information, such optimized workflows often feature a low number of MS1 scans, frequently acquired at resolutions below 60k. Consequently, the majority of publicly available datasets are suboptimal for search engines that require MS1 isotopic envelopes profiles as the input information, such as FragPipe-DIA or msallsearchpy. A notable example is the dataset from Kawashima et al.^24^, for which DIA-NN reports 98,000 peptides and 10,600 protein groups at 1% FDR, whereas FragPipe-DIA identifies only 37,000 peptides and 7,900 protein groups. This dramatic difference is largely explained by a distinct feature of the data: approximately 50% of the identifications reported by DIA-NN lack confirmation from precursor ion isotopic envelopes. This lack is likely a consequence of the low MS1 resolution of only 15K employed for the MS1 spectra acquisition. Because the original work aimed specifically to maximize protein identifications via DIA-NN, minimizing the instrument time allocated to MS1 spectra was a logical strategy to increase those specific target metrics, albeit at the expense of MS1 data quality.

Unlike the standard DDA/DIA search engines which are MS/MS-centered, msallsearchpy primarily searches MS1 spectra for identifications. As demonstrated elsewhere, MS1-based quantification represents the strategy of choice for high-throughput proteomics^25^. With this regard, msallsearchpy is an peptide isotopic envelope-centric search algorithm that unifies MS1-only, DDA and DIA data analyses. The primary limitation of this algorithm is its absolute dependence on the quality of the MS1 spectra. In the absence of discernible peptide precursor signals at the MS1 level, the algorithm cannot yield a meaningful number of identifications, regardless of the abundance of high-quality MS/MS spectra. The core paradigm underlying this strategy is the deliberate exclusion of any peptide identifications that lack confirmation from high-quality MS1 isotopic envelopes. Among the existing DIA algorithms, a notable example is FragPipe-DIA, which also requires the presence of precursor isotopic envelopes in MS1 spectra for identified peptides. However, a key distinction remains: FragPipe-DIA utilizes a targeted approach to extract peptide isotopic envelopes, whereas msallsearchpy analyses raw MS1 spectra independently of prior database knowledge via untargeted feature detection. The latter paradigm is rarely deployed in the field with a few examples, such as DIA-Umpire^26^ and Quandenser^27^. Conversely, certain DIA algorithms do not require the presence of an MS1 precursor isotopic envelope to assign peptide identities, rendering them incapable of distinguishing between peptides that share highly similar y-series fragments^28^. Although, this limitation might initially appear to affect only peptide-centric applications, such as proteogenomics, immunopeptidomics, or post-translational modification (PTM) analysis, the presence of the non-random, biologically conserved amino acid motifs introduces a substantial risk to the results. Specifically, ignoring precursor information increases the likelihood of falsely identifying incorrect proteins that share sequence motifs with the true components present in the sample. Further, we argue that quantitative benchmarks obtained using multi-species proteomes may be compromised by false matches, as shared amino acid motifs following the fold change distribution of the true peptides from the same species. This specific vulnerability underscores our rationale for selecting FragPipe as our benchmarking baseline among the available range of commonly used workflows, as its architecture ensures data reliability.

A current limitation of msallsearchpy is its lack of focus on peptide-level identifications, including the absence of support for variable modifications. Thus, the search engine is not currently suited for peptide-centric applications, such as proteogenomics, PTM profiling, degradomics, etc. However, conventional proteomic studies aimed at efficient detection of differentially expressed proteins, will heavily benefit from the msallsearchpy workflow.

In summary, the msallsearchpy engine offers a unique combination of features, including an open source license, broad universality, and the utilization of the standard, well-characterized target-decoy approach for FDR control. The software is compatible with practically all major types of acquisition methods, including MS1-only (DirectMS1), DDA, and DIA. This universality was achieved by integrating available MS/MS information directly into the core MS1-based search framework. The primary advantage of msallsearchpy is its exceptional efficiency when processing data acquired using micro- and analytical-flow regimes on Orbitrap mass spectrometers, particularly in the currently mainstream analyses of low sample amounts and short LC gradients.

## Online Methods

A description of all datasets used in the study is shown in Table 1. A brief description of all raw files is provided in Supplementary Table S1.

**Table 1.** Datasets used in the study.

| ProteomeX change ID | Figure | Flow rate, ul/min | Acquisition | Sample | Instrument | Sample amount, ng | Gradient, min | Type of data |
| --- | --- | --- | --- | --- | --- | --- | --- | --- |
| PXD026600 | S1; S2 | Nano | DIA | *E. coli* | Fusion | 1000 | 120 | ID-only |
| PXD062536 | 2A; S3; S4 | 1.5-50 | DDA | HeLa | QE HF-X | 20-2000 | 60 | ID-only |
| PXD057525 | 2B; S5 | 115; 51 | DDA | Jurkat | QE HF | 50-5000 | 15;60 | ID-only |
| PXD069554 | 2C | 50 | DDA | Jurkat | QE HF-X | 160-16000 | 60 | ID-only |
| PXDXXXXX | 3a | 400 | DIA | A549 treated with bortezomib | QE HF | 500; 5000 | 25 | Quant |
| PXDXXXXX | 3b | 400 | DIA | PISA, HCT116, staurosporine | QE HF | 500 | 25 | Quant |
| PXD050784 | 3b | 0.25 | DIA | PISA, staurosporine | Lumos | Unknown | 50 | Quant |
| PXD070049 | 4;S6 | Nano | DIA; DDA | K562/yeast/*E. coli* “LFQ Bench” | Astral | 0.25 | 15; 5 | Quant |

### Datasets

Raw DDA data from Tsiklauri et al.^10^ was downloaded from ProteomeXchange^29^ (PXD062536). We re-analysed the data for 20, 200 and 2000 ng sample loading amounts. Flow rates and inner column diameters were: 50 ul/min and 1 mm, 10 ul/min and 300 um, 5 ul/min and 300 um, and 1.5 ul/min and 150um.

Raw DDA data from adeja et al.^13^ was downloaded from ProteomeXchange (PXD057525). We re-analysed the 15 and 60 min gradient data for 1.0 mm i.d. (51 ul/min) and 1.5 mm i.d. (115 ul/min) columns.

Raw DDA data from Starovoit et al.^9^ was downloaded from ProteomeXchange (PXD069554). We re-analysed the 60 min gradient data for propionic and formic acids.

Raw DDA and DIA data from Puyvelde et al.^30^ was downloaded from ProteomeXchange (PXD070049). We re-analysed the 5 and 15 min Orbitrap Astral data for samples containing a mix of human, yeast and *E. coli* proteins. Conditions A and B were used where experimental log2 fold changes between conditions for yeast and *E. coli* proteins were -1 and 2, respectively. Human proteins were not changed between conditions. We analysed only the data with a total sample loading amount of 250 pg which was generated to emulate “single-cell equivalent” inputs.

A single raw file containing narrow-window DIA data for 2.5 fmol of UPS proteins spiked into 1 ug of *E. coli* proteins was downloaded from ProteomeXchange (PXD026600).

Newly generated analytical flow DIA data for expression proteomics with Bortezomib treatment and PISA TPP proteomics with Staurosporine treatment were uploaded to ProteomeXchange (PXDXXXXX).

Raw DIA data from Batth et al.^22^ was downloaded from ProteomeXchange (PXD050784). We re-analysed the data for proteins with centrifugation protocol analysed using 50 min LC gradient, Orbitrap Lumos, and 250 nL/min flow rate. The temperature sample pooling for PISA was performed using three temperature points: 50, 53, and 56 °C.

### Sample preparation

#### PISA TPP

A total of 40×10^6^ HCT116 cells were resuspended in 600 µL of lysis buffer containing 240 µL of protease inhibitor cocktail (cOmplete™, EDTA-free from Roche, Switzerland) and 360 µL of 1× PBS solution. The suspension was divided into three aliquots, and cells were lysed by five freeze–thaw cycles, each consisting of 2 min in liquid nitrogen followed by 2 min in a 37 °C water bath. Lysates were centrifuged at 20,000g for 20 min at 4 °C. The supernatants were collected and pooled, and protein concentration was measured by a BCA Protein Quantitative Detection Kit (ServicуBio, China).

The stock solution was divided into control and treatment samples, and PBS solution was added to reach the final protein concentration of 1.5 µg/µL. Staurosporine was prepared as a 1 mM stock solution in DMSO and added to the treatment group at a 1:100 dilution to obtain a final concentration of 10 µM. Control samples received an equal volume of DMSO at the same final dilution (1:100, v/v).

Samples were incubated at 25 °C for 35 min, followed by extraction of the supernatant and determination of protein concentration as described above. Then, both treatment and control samples were divided into seven temperature groups, with four technical replicates per temperature point. Samples were heated for 3 min at 40, 44, 48, 52, 56, 60, or 64 °C and immediately cooled on ice.

Following heat treatment, aliquots corresponding to all temperature points were pooled within each experimental condition to generate pooled samples. In parallel, an additional set of 40°C samples was processed separately. Both the pooled samples and the separately prepared 40 °C samples were centrifuged at 50,000 g for 30 min at 4 °C. Supernatants were harvested and DTT (10 mM) was added to samples for 20 minutes at 56°C. Then the samples were incubated with IAA (15 mM) for 30 minutes in the dark at room temperature.

Proteins were subsequently precipitated using a chloroform–methanol precipitation procedure. Protein pellets were resuspended in 40 µL of 0.1% ProteaseMAX surfactant (Promega, USA) in 100 mM ammonium bicarbonate (ABC) and 40 µL of 100 mM ABC. Samples were vortexed and sonicated in an ultrasonic bath for 5 min. Trypsin (Trypsin LC-MS grade, Molecta, Russia) was added with a ratio of 1:75 (w/w), and samples were incubated overnight at 37°C. Digestion was stopped by adding 1% TFA, and then samples were centrifuged for 15 min at 20,000g; supernatants were taken and dried in a vacuum concentrator.

#### Expression proteomics

Samples prepared for Vorobeva et al.^31^ were reused. Briefly, A549 cells were treated with Bortezomib dissolved in the culture medium containing 0.05% DMSO for 48 hours with IC_50_ concentration (0.3 uM). Control samples were treated using the same medium with DMSO. Trypsin digestion was used to prepare samples for LС/MS analysis.

#### DIA LC-MS/MS

MS analysis of proteomic samples in this work was performed using Orbitrap Q Exactive HF mass spectrometer (Thermo Scientific, USA) coupled with UltiMate 3000 LC system (Thermo Fisher Scientific, Germany). Luna column (C18; 100 Å, 5 µm; 2.0 mm × 150 mm) was employed for separation. Mobile phases were as follows: (A) 0.1 % formic acid (FA) in water; (B) 80 % ACN, 0.1 % FA in water. The gradient was from 5 % to 35 % phase B in 20 min at 400 µL/min. Total method time including column washing and equilibration was 34 min. Mass spectrometry measurements were performed in DIA mode of acquisition. Full MS scans were acquired in a range from m/z 375 to 1200 at a resolution of 120 000 at m/z 200 with Automatic Gain Control (AGC) target of 3·10^6^ and 1 microscan. Maximum injection times were 50 and 100 ms for expression and TPP proteomics, respectively. Full cycle consisted of 4 MS1 and 28 MS2 scans in the precursor m/z range of 462–840 with a 14 Da step, an isolation window of ±7.5 Da (i.e., with slight overlap), and a resolution of 30k. MS1 scan was acquired after every 7 MS2 scans to increase the coverage of peptide retention time profiles by MS1 scans.

### Data processing

Raw files were converted to mzML format using ThermoRawFileParser^32^ (v. 1.4.5).

#### msallsearchpy

Peptide isotopic envelopes were extracted from MS1 spectra using biosaur2 (v. 0.3.2). Parameters for the biosaur2 were as follows: charge states from 2+ to 6+,, minimum 2 scans for detected isotopic cluster, 8 ppm mass tolerance for hill profiling and 8 ppm mass tolerance for isotope envelope construction. The minimal number of consecutive MS1 scans was set as 1 for Orbitrap Astral DIA 5 min and our Orbitrap QE DIA data. ms1searchpy (v. 3.0.2) was used to process DDA/DIA data. Parameters for the search were as follows: “msallsearchpy mode” (to extract MS/MS spectra information), DeepLC^17^ (v. 4.5) to predict peptide retention times, 12 ppm initial precursor mass tolerance, 1% identification protein-level FDR, 7-30 amino acid length for fully-tryptic peptides, carbamidomethyl of cysteine as fixed modification. Precursor mass tolerance and compensation of systematic mass shift were set as 6 ppm and 5 ppm, respectively, for Orbitrap Astral DDA data.

DirectMS1Quant^3^ was used for differential expression analysis using default settings for all datasets: 50% maximum missing values for peptides, median intensity normalization by retention time intervals, 2.5 standard deviations of background protein fold changes as fold change threshold, 5% quantitative FDR. Keratin proteins were removed from the results for Staurosporine data.

#### Identipy

IdentiPy^20^ (v. 0.3.9.5) was used to analyse high-flow DDA data. The search options were: 10 ppm and 0.03 Da precursor and fragment mass tolerances, respectively, up to 1 isotope precursor selection error, 1 allowed missed cleavage. Postsearch analysis of pepXML files was performed using Scavager^33^ (v. 0.2.18) with 1% FDR settings.

#### FragPipe

FragPipe (v. 24) was used as a representative of state-of-art proteomics software. All search options were default ones for workflows “DIA_SpecLibQuant” and “LFQ-MBR” for DIA and DDA data, respectively. The only difference from the standard workflows is the shuffled decoy database generated by msallsearchpy was used instead of the default reversed one. FragPipe workflow integrated multiple algorithms, including MSFragger^34^ (v. 4.4.1), MSBooster^35^ (v. 1.4.14), Percolator^36^ (v. 3.7.1), Philosopher^37^ (v. 5.1.3-RC9), IonQuant^38^ (v. 1.11.20) and DIA-NN^39^ (v. 1.8.2 beta 8). FragPipeAnalyst (v. 1.26) was used to extract differentially expressed proteins after FragPipe processing. For all data, the options were 0.05 adjusted p-value cutoff, Benjamini-Hochberg correction, 0.5 log2 fold change cutoff, Perseus-type imputation, no normalization. For DDA data, the “MaxLFQ Intensity” option was used. An option “Remove contaminant proteins” was enabled for Staurosporine data.

### msallsearchpy algorithm

#### Protein identification workflow

The core algorithms of the workflow were kept unchanged since our previous publications^3,5,6^. A modified pseudo shuffle decoy generation method is used for the FDR control, which can deal with the databases containing proteins with high homology (details are provided in the supporting information of the directms1quant study^3^). Protein scoring is performed using a binomial model, where random peptide matching probabilities are estimated using the decoy matches. LightGBM gradient boosting is used to estimate the reliability of peptide matches, but the peptide-level FDR is not strictly controlled.

#### MS/MS scoring

MS/MS spectra are used without any clustering and demultiplexing stages for both DDA and DIA data in msallsearchpy. Every applicable (by isolation window) experimental MS/MS is scored against the theoretical fragmentation spectra of peptide sequence candidates. Three types of search scores are calculated: a default RNHS score which was introduced in IdentiPy search engine, a score adjusted for better chimeric spectra processing, and a probability-based score. The former is close to X!Tandem hyperscore, which is calculated as multiplication of factorials of matched b and y ions, as well as the fraction of matched fragment intensities. The second one is different by the intensities-based multiplicator - the fraction of matched fragment intensities is normalized by the sum of intensity values less than the maximal intensity of the matched fragments instead of the sum of intensities of all experimental fragments. The idea is that chimeric fragments may have extremely low intensity values and there be a discrimination for such matches in the standard scores. The third score is calculated as the binomial distribution survival function (SF(k, n, p)), where probability p is calculated as the average of the individual fragment ions probabilities to be matched randomly. These individual probabilities are estimated using all matched decoy peptides of the same length. The k and n are the numbers of matched and total theoretical fragments. The spectrum with maximum RNHS score is detected first for the particular peptide and the rest of scores are calculated for that particular spectra. This is done to optimize the processing time. A fourth score (optional) is calculated if the MS2PIP option is enabled. The score is a Pearson correlation between experimental fragment intensities and the ones predicted by MS2PIP. All these MS/MS-based scores, as well as the number of matched b and y ions are added to the peptide-feature match information which is used in the machine learning-based classification done in the standard realization of ms1searchpy algorithm. Thus, the presented search workflow is close to the MS1-only algorithms (DirectMS1), where some of the matched PFMs are extended with useful MS/MS-based information. It means that the PFMs with no matched fragments are still fully utilized in the protein identification and quantification processes.

#### Retention time prediction with DeepLC

The original ms1searchpy implementation utilized DeepLC of version 0.1.2. Starting from ms1searchpy v.3.0.2 (the one applied in this study) the utilized DeepLC version was upgraded to 4.5.0. Note that we did not use the calibration algorithm built into the DeepLC package, as the latter does not perform well in our case due to the presence of false identifications in the calibration set of peptides. A custom model selection procedure among the available 6500 pre-trained models was implemented within ms1searchpy. Up to 2500 most reliable peptide matches are selected and split into two random equal groups. The first group is used to calibrate pre-trained DeepLC model for experimental RTs using piecewise linear regression and the second one is used to estimate standard deviation of retention time prediction error by fitting retention time differences with the sum of normal (true peptide matches) and uniform (false peptide matches) distributions. Next, the steps are repeated by swapping the peptide groups. The worst (highest) standard deviation of RT error is saved for the particular model and all the steps are repeated for the next model (DeepLC of version 4.5.0 provides ∼6500 pre-trained models). Finally, the model with the best (lowest) std of RT error is used in the algorithm for all peptides within the particular experimental run.

#### Differential expression analysis using DirectMS1Quant

Quantification is based on the peptide’s monoisotopic peak apex intensities. Differential expression analysis is described elsewhere^3^ with three key updates introduced in this work. First, we changed the intensity normalization process. In its original realization, peptide’s intensities were normalized by median intensities of quantified peptides in each analysed file. Now, the intensities are normalized by median from 50 uniformly distributed intervals of retention times for quantified peptides. The algorithm is our reimplementation of the one described in the Lyutvinskiy et al.^40^

Second, we changed the p-value calculation of differential expression. In its original realization, the p-value was calculated using the survival function of binomial distribution, where k is the number of significantly expressed peptides (for particular protein), n is the number of identified peptides (for particular protein) and p is the probability of peptide to be significantly expressed by random. The latter is estimated using quantification of decoy PFMs and proteins. Now, we extended the results with a second type of p-value, which is close to the traditional differential expression analysis, where p-value is calculated using an independent t-test with protein LFQ values used as input. Those two p-values are combined into the final one using Fisher’s combined probability test.

Thirdly, quantified peptides are now matched between files by using retention times. For example, if some peptide was identified in all experimental files, but the retention times are 50, 49, 51, 52, 48 and 90 minutes, the last one will be excluded from the analysis. Allowed retention time threshold is determined automatically. Peptides identified in all files are selected and the difference between the particular value of retention time and the median value of retention times for these peptides is calculated. Next, values for 10% and 90% percentiles of those differences for all peptides are calculated. And finally, doubled values of those percentiles are used as the allowed retention time difference between files for all quantified peptides.

#### Speed processing optimization

The idea to optimize speed of fragments matching process by using integer approximation of precursor and fragment masses is not novel and quite broadly adapted in proteomics^34,41,42^. For years we used a similar approach in our time-extensive algorithms like IdentiPy DDA search engines or biosaur2 peptide feature detection, but that update was not done in the original builds of the corresponding software and thus was not described in the manuscripts. While there is no strong novelty, here we provide brief details of our implementation for the first time which may be useful for the developers in the proteomics field. Unlike the common solutions, which use the sparse matrix which significantly reduces the processing time at the cost of RAM memory, we use a dictionary structure (called dict in Python, unordered_map in C++ and HashMap in Java programming languages) which is compact. All precursor and fragment masses are divided by used-defined precursor and fragment mass accuracies recalculated in Da values for the maximal experimental m/z values of precursors and fragments. The resulting values are rounded to the integer type. This is done for both theoretical and experimental spectra. The theoretical spectra is then stored in a “set” Python structure, while the experimental spectra is stored in a “dictionary” structure with integer masses stored as the keys and intensities normalized by the sum of intensities for the spectra are stored as the values. Every experimental integer mass M is tripled as M-1, M and M+1 values. As a result, the simple intersection operation (time complexity is O(1)) between set and dictionary key results in a list of intensities, as well as the number of matched fragments. When we use those obtained values, we can calculate upper approximation for RNHS score discussed above. Only if this upper approximation is higher than the current best RNHS score for a particular peptide, we calculate an accurate RNHS score. Because set intersection operation is extremely fast, we can significantly reduce the processing time and don’t extend RAM usage, while keeping our Python-based algorithms competitive to the other proteomics algorithms developed in fast and optimal languages like C, Java or Rust.

#### Runtime

The full msallseachpy quantitative workflow is based on three software tools: biosaur2 for peptide isotopic ion envelopes detection, ms1searchpy for protein identification and directms1quant for differential expression analysis. The code is based on the Python programming language which makes multiprocessing utilization a non-trivial task. Because of that, the number of processes used by default on Windows-based computers is limited to one. However, unlike the classic DDA/DIA search engines, msallsearchpy does not have any match-between-runs information shared during the identification stage. Thus, the biosaur2 and ms1searchpy commands can be executed in parallel for each file as soon as there is sufficient volume of available RAM to process the entire dataset of interest. directms1quant is the only command which cannot be executed in parallel. The runtime per randomly selected file from 15 min LC DDA Astral dataset were 0.5 and 6 min for biosaur2 and ms1searchpy, respectively (For 2 X EPYC 7742, 1 Tb RAM and RTX A4000 PC). Both commands were executed with an option to use only one CPU core. Note, that the number of cores available for the DeepLC was not controlled. Thus, the total runtime using only one truly available core was estimated as 23 min instead of 6. Runtime of directms1quant per one condition (4 replicates vs 4 replicates) was 2 min. Peak RAM consumption were 1, 4.5 and 0.4 Gb for biosaur2, ms1searchpy and directms1quant, respectively. Peak GPU RAM consumption (utilized by DeepLC) was 0.5 Gb. All 12 DDA Astral files were processed in parallel in 1 and 6 min for biosaur2 and ms1searchpy, respectively. No significant drops in execution time per file were found when the 7 years old PC was used (i9-9900X, 128 Gb RAM and RTX 2060 PC): 0.3 and 7 min for biosaur2 and ms1searchpy, respectively. There are plenty of possibilities to speed up calculations and RAM consumption in the future updates, but the current performance is reasonable for most user cases and it is out of the focus of our current developments.

## Code Availability

The Jupyter notebook and all the search files used to generate Figures 2–4 and the Supplementary Figures 1-7 are available through Zenodo, under 10.5281/zenodo.23321052. The msallsearchpy search engine is available at https://github.com/markmipt/ms1searchpy under Apache 2.0 license. The repository contains a quick guide on how to reproduce msallsearchpy’s output files for “LFQ bench” dataset in just five terminal commands on a Linux system. Other results from this work were obtained in a similar manner, with minor adjustments to the parameters clarified in the methods section.

## AUTHOR CONTRIBUTIONS

M.V.I. conceived and supervised the project, developed the algorithm, wrote the software, and analysed the results. L.A.G. and V.I.P. contributed to software development, testing and data analysis. E.M.K. contributed to software testing. L.A.G. made biological experiments and sample preparation. A.A.S. made LC/MS analysis. D.D.E., E.M.K., and I.A.T. contributed to proteomics experiments. A.V.T. contributed to LC/MS analysis. M.V.G. provided critical feedback and discussed the results. M.V.I. and M.V.G. wrote the manuscript.

## Notes

The authors declare no competing financial interest.

## ACKNOWLEDGMENTS

The study was performed with financial support from the Russian Science Foundation, grant no. 25-74-10042 to M.V.I.

---

**Supplementary Figure S1.**
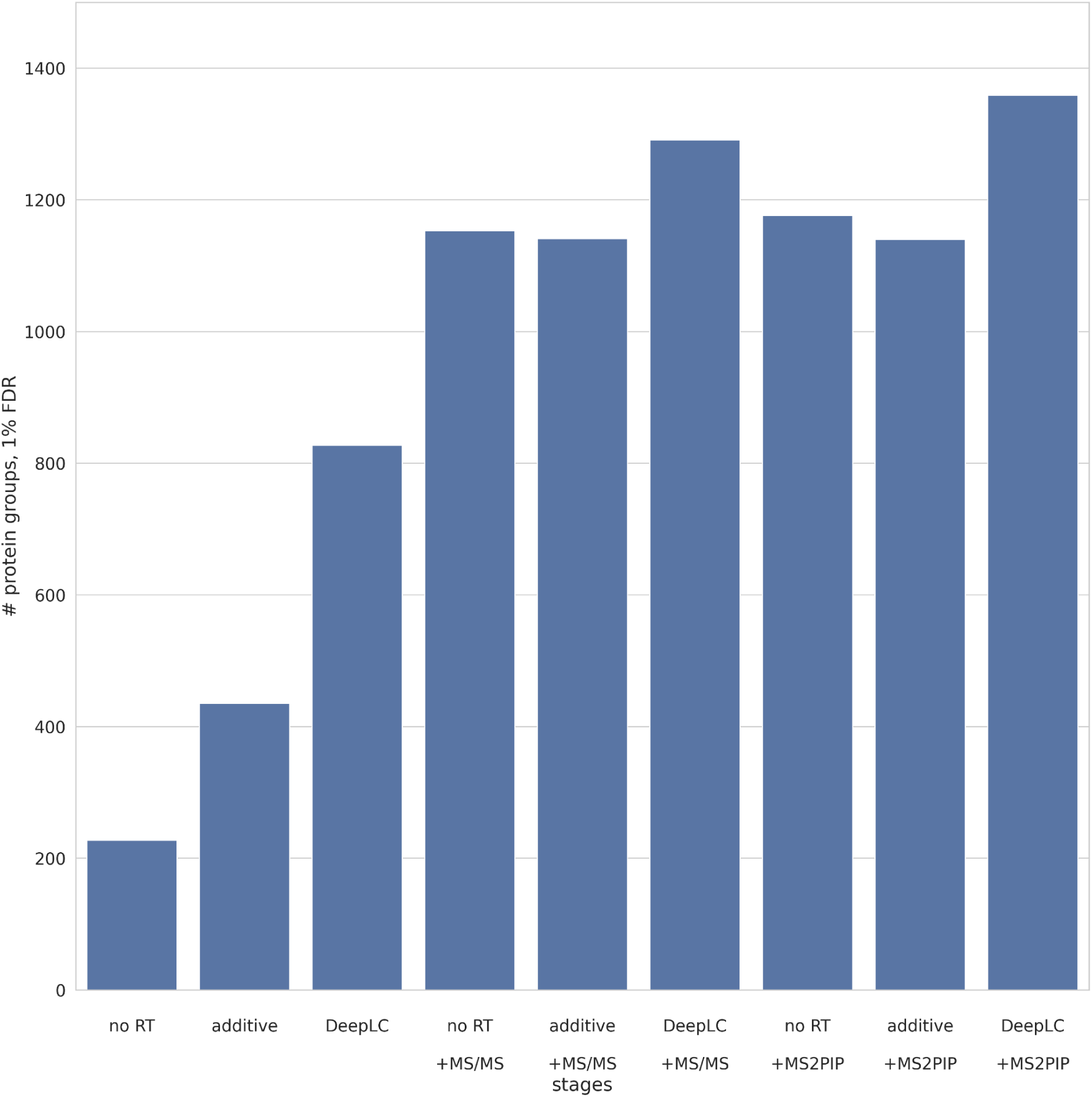
Influence of different data processing stages on the number of identified proteins. The results are shown for the UPS-*E. coli* PXD026600 dataset.

**Supplementary Figure S2.**
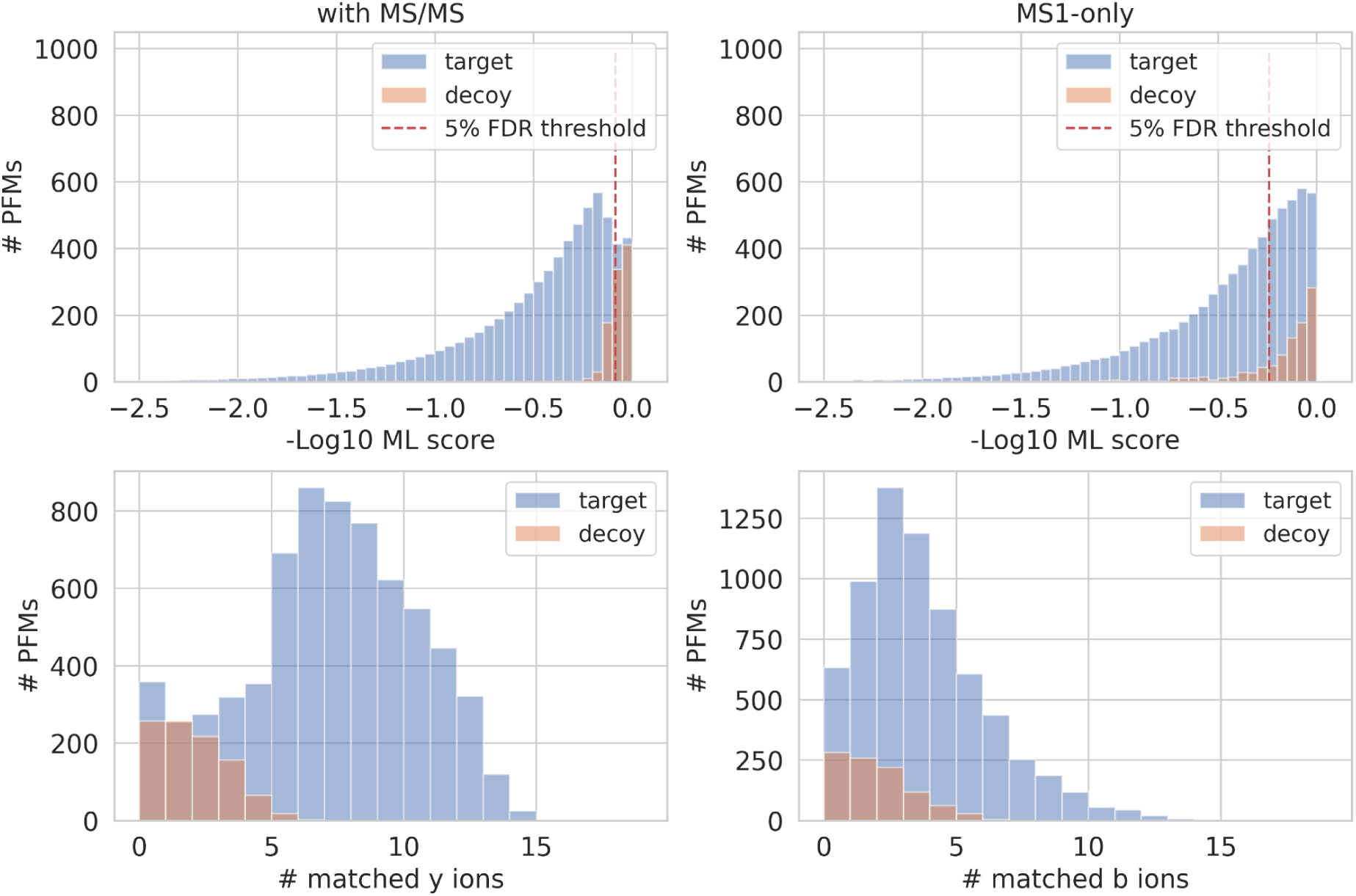
Distribution of matched target and decoy PFMs by machine learning scores and the number of matched fragment ions. The results are shown for UPS-*E. coli* PXD026600 dataset.

**Supplementary Figure S3.**
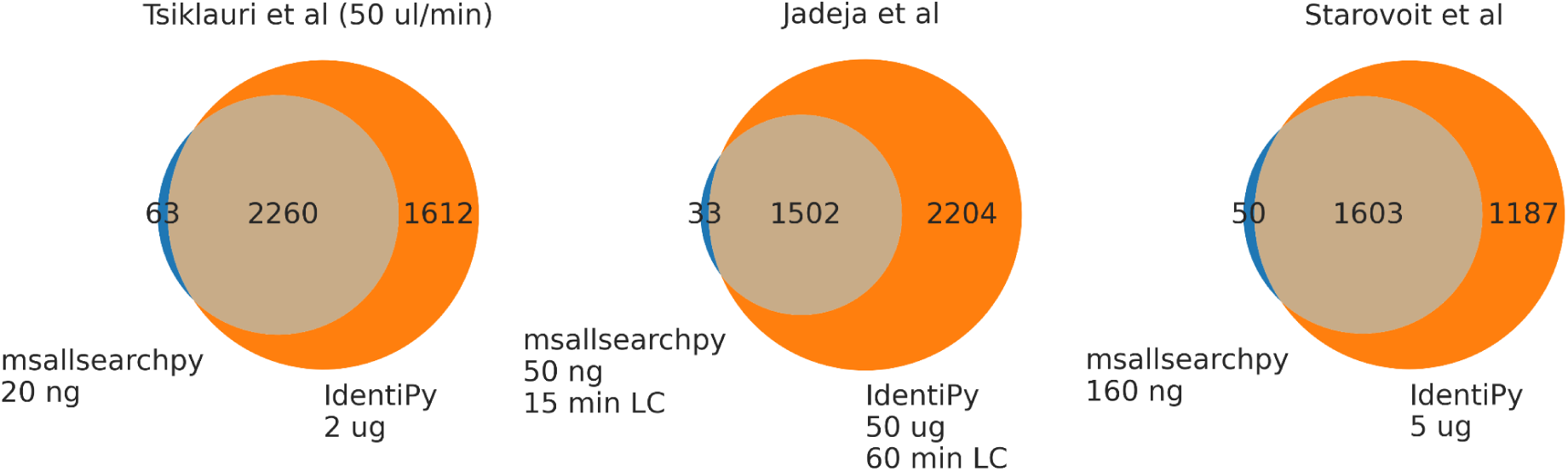
Venn diagrams for the identified protein groups by msallsearchpy and IdentiPy for selected conditions and datasets. Single random replicate per condition was used.

**Supplementary Figure S4.**
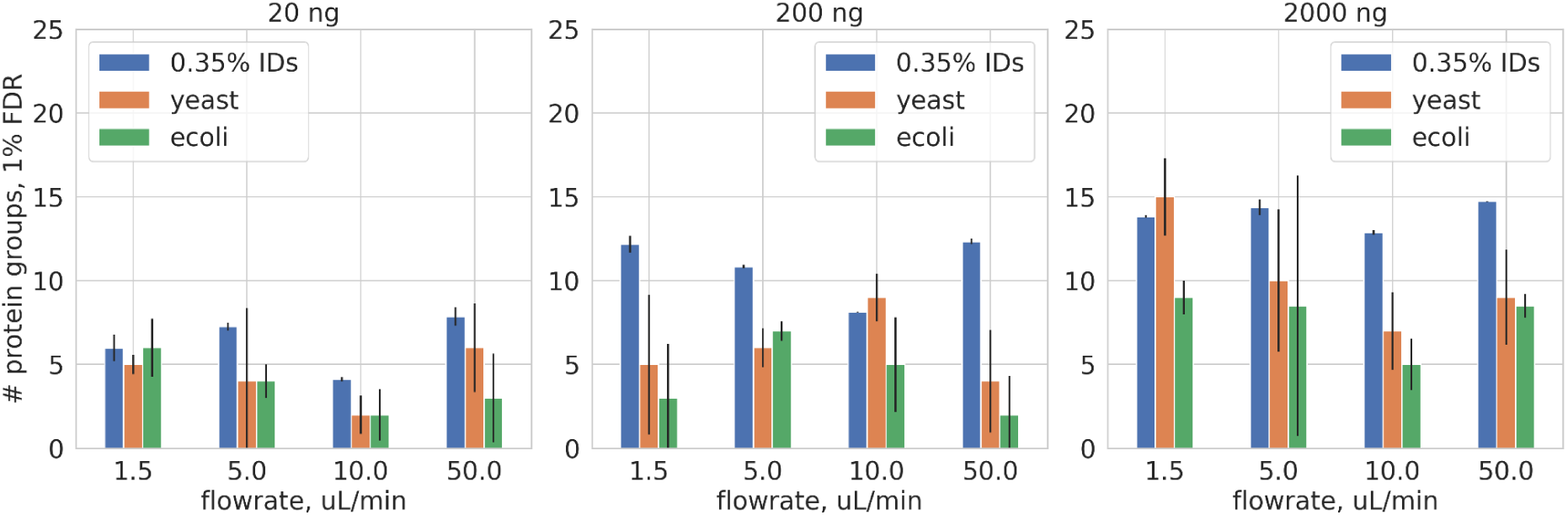
The number of identified *E. coli* and yeast protein groups by msallsearchpy at 1% FDR for Tsiklauri et al. when combined human + *E. coli* + yeast protein database was used. The blue color represents the 0.35% of total number of identified protein groups, which is the expected number of *E. coli* and yeast false identifications according to the relative database sizes.

**Supplementary Figure S5.**
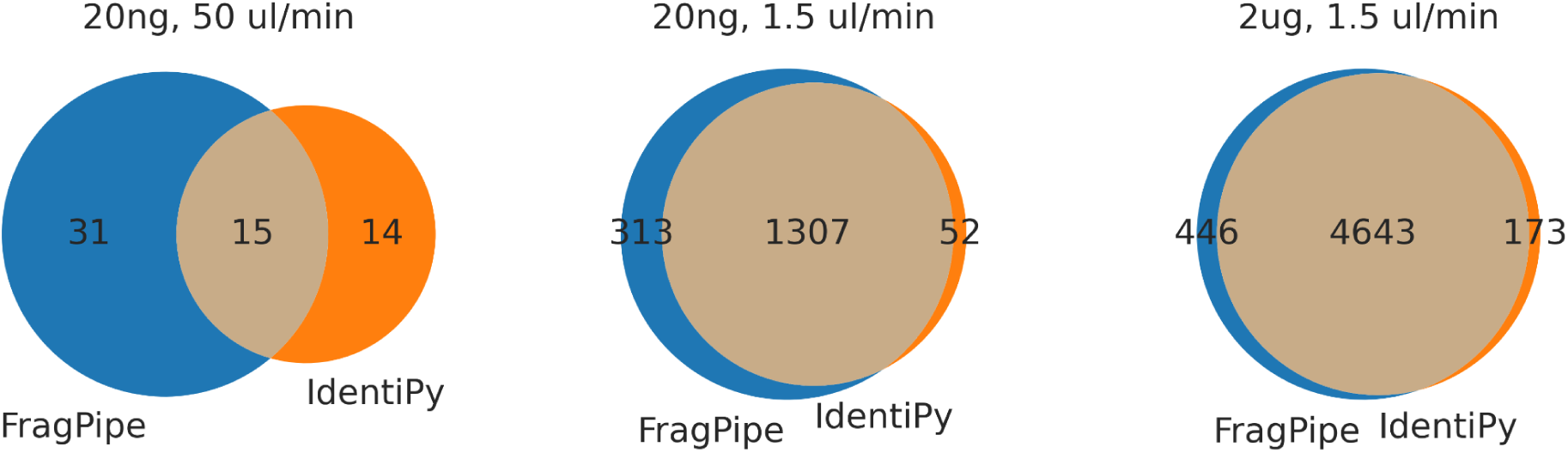
Venn diagrams for the identified protein groups by FragPipe and IdentiPy for selected conditions dataset from Tsiklauri et al. Single random replicate per condition was used.

**Supplementary Figure S6.**
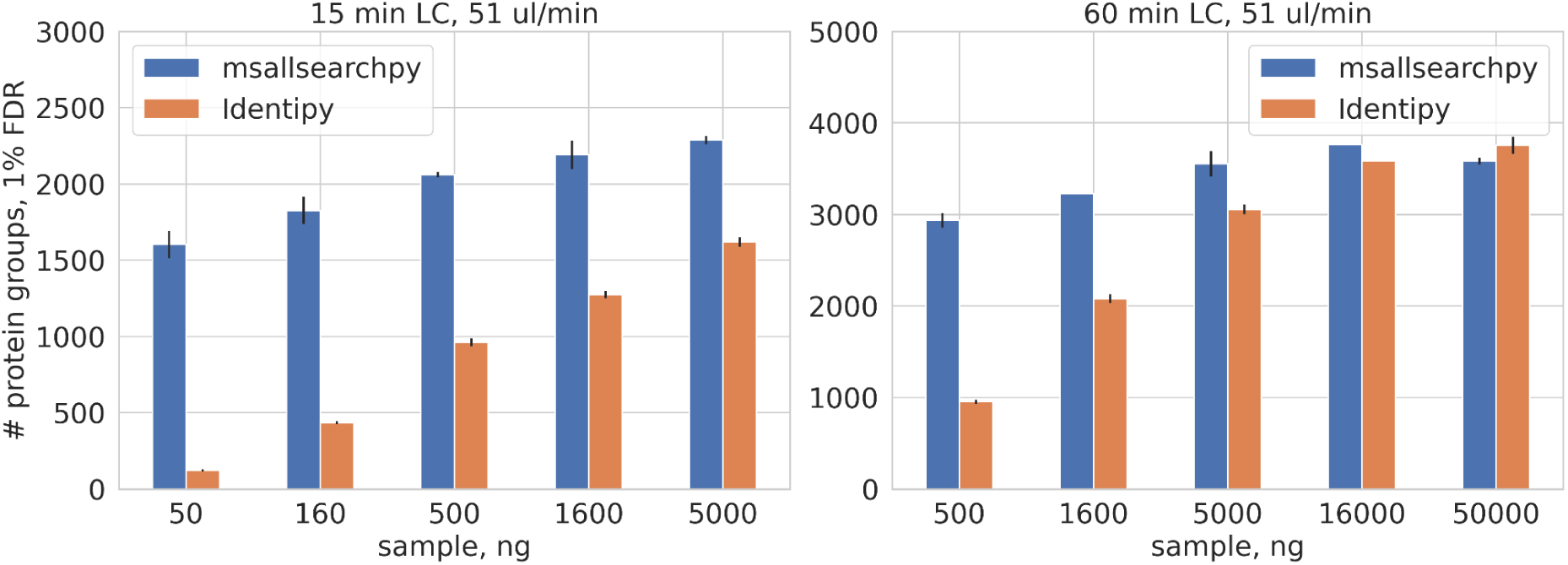
The number of identified protein groups at 1% FDR for 1.0 mm i.d. LC column data from Jadeja et al. Results for msallsearchpy are indicated in blue, while results for the traditional DDA search engine, Idenipy, are shown in orange.

**Supplementary Figure S7.**
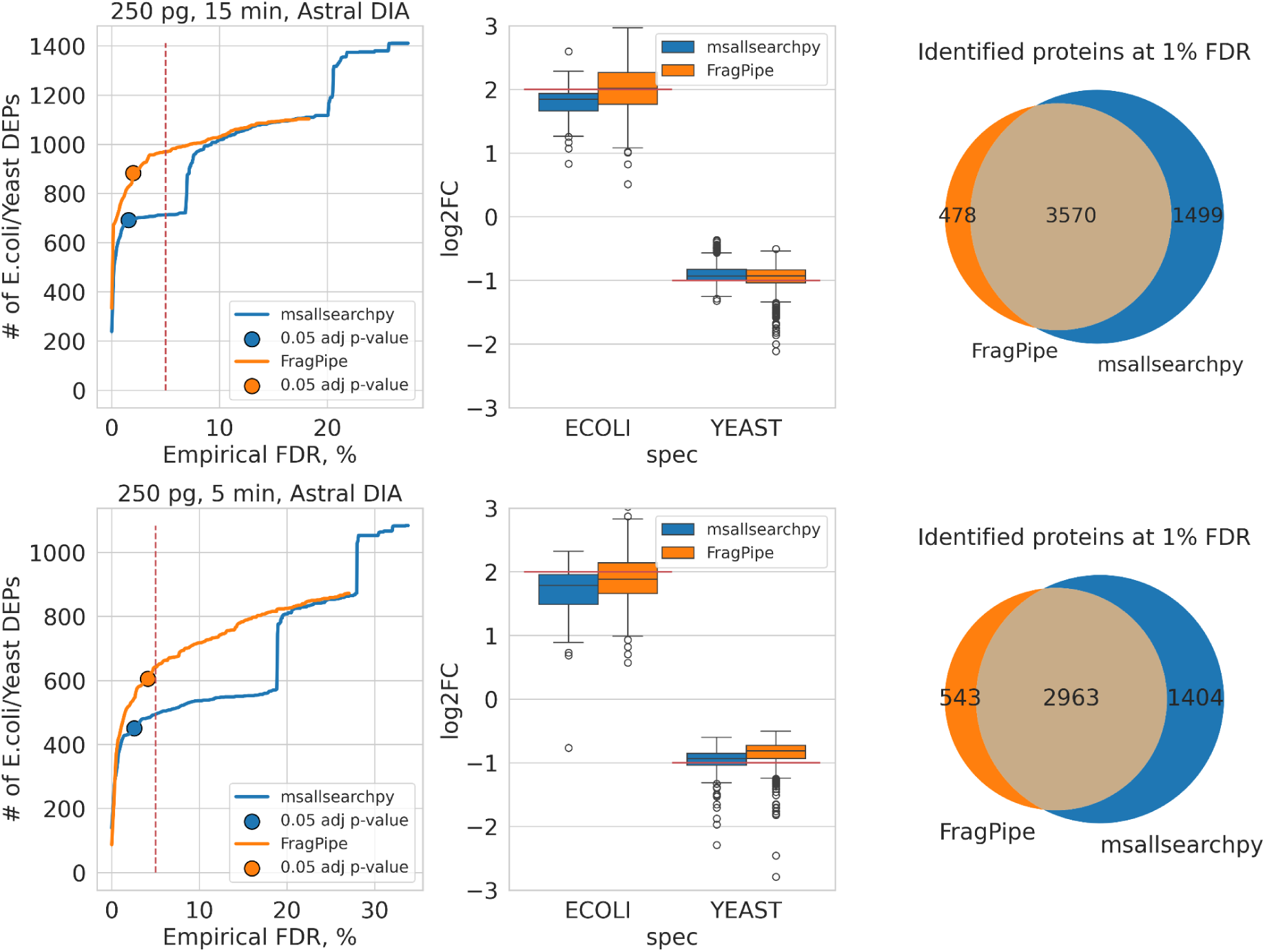
Results for 250 pg Orbitrap Astral DIA data from Puyvelde et al. analysed by msallsearchpy+DirectMS1Quant (blue) and FragPipe+FragPipeAnalyst (orange) workflows: (upper figures) 15 min LC and (bottom figures) 5 min LC data. (left column) The number of reported differentially expressed *E. coli* and yeast proteins. Empirical FDR was calculated using reported human DEPs. Only the proteins that passed fold change thresholds were used for the plots. Fold change thresholds were: 0.5 absolute log2 value for FragPipeAnalyst and automatic for DirectMS1Quant. Blue and orange dots represent the results which were obtained using 0.05 p-value threshold with Benjamini-Hochberg correction with no prior knowledge on the empirical FDR. *E. coli* and yeast proteins with opposite to the real changes sign of log2 fold changes were considered as false DEPs. (middle column) The log2 fold changes of DEPs. Red lines represent the experimental fold changes. (right column) The number of identified protein groups at 1% FDR.

## Notes

### Competing Interest Statement

The authors have declared no competing interest.

